# Autophagy dark genes: Can we find them with machine learning?

**DOI:** 10.1101/715037

**Authors:** Tudor I. Oprea, Jeremy J. Yang, Daniel R. Byrd, Vojo Deretic

## Abstract

Identifying novel genes associated with autophagy (ATG) in man remains an important task for gaining complete understanding on this fundamental physiological process. A machine-learning guided approach can highlight potentially “missing pieces” linking core autophagy genes with understudied, “dark” genes that can help us gain deeper insight into these processes. In this study, we used a set of 103 (out of 288 genes from the Autophagy Database, ATGdb), based on the presence of ATG-associated terms annotated from 3 secondary sources: GO (gene ontology), KEGG pathway and UniProt keywords, respectively. We regarded these as additional confirmation for their importance in ATG. As negative labels, we used the OMIM list of genes associated with monogenic diseases (after excluding the 288 ATG-associated genes). Data associated with these genes from 17 different public sources were compiled and used to derive a Meta Path/XGBoost (MPxgb) machine learning model trained to distinguish ATG and non-ATG genes (10-fold cross-validated, 100-times randomized models, median AUC = 0.994 +/− 0.0084). Sixteen ATG-relevant variables explain 64% of the total model gain, and 23% of the top 251 predicted genes are annotated in ATGdb. Another 15 genes have potential ATG associations, whereas 193 do not. We suggest that some of these 193 genes may represent “autophagy dark genes”, and argue that machine learning can be used to guide autophagy research in order to gain a more complete functional and pathway annotation of this complex process.

## Introduction

Autophagy (ATG) is a cytoplasmic homeostatic process defined by a suite of ATG genes conserved from yeast to man [1]. Autophagy keeps a cellular complement of organelles in a functional state and removes protein aggregates, invading pathogens, and other cargo, by capturing them and delivering them to lysosomes for degradation [1]. Autophagy is consequential for a wide array of physiological and pathological states including neurodegenerative and other degenerative disorders, cancer, chronic inflammatory illnesses and infectious diseases [2]. Mechanistically, the process of autophagy is governed by ATG genes, encoding the core autophagy machinery with most components conserved from fungi to humans [1]. However, recent studies have increasingly emphasized the existence of systems controlling and executing autophagy in mammalian cells that are quite different from those in yeast [3]. Among well-accepted examples, are several autophagy factors absent in yeast that have been identified in organisms from *C. elegans* to *H. sapiens,* including *ATG101* [4], *FIP200* [5], *VMP1* [6], *EPG5* [7], *Stx17* [8,9], and *TMEM41B* [10–12].

Completing the autophagy puzzle requires the systematic identification of all genes associated with autophagy. This on-going endeavor is boosted in part by the growing interest in the role of autophagy in ageing regulation [13,14] and lifespan extension [15]. ATG genes have been used in supervised machine learning models applied to ageing research [16] and are among the top features in models for predicting lifespan-extending chemicals [17]. Numerous studies have used machine learning (ML) methods to infer gene-disease associations [18–21]. However, the use of ML models to guide further autophagy research has not been previously discussed. Here we report the development of a specific ML model to predict “autophagy dark genes”, i.e., understudied proteins [22] that may play a significant role in autophagy.

## Materials and Methods

### Building Knowledge Graphs

Seventeen different data sources, totaling over 262.3 million data points, summarized in **Table 1**, are used to build Knowledge Graphs (KGs). These 17 protein-and gene-centric data sources are integrated into formal data representation systems based on **KGs**, with typed nodes and edges, which enable the use of network-based analytical algorithms. Whereas most of the data in Table 1 refer to human proteins, we use gene orthology relationships from eggNOG [23] and InParanoid [24] to fuse rat (RGD) and mouse (IMPC, International Mouse Phenotype Consortium) model organism data into a single “pseudoprotein” that enables network-based inferences for function and phenotype across organisms. The **KGs** are implemented as a PostgreSQL db for performance, convenience, and integration with existing tools. Metapaths are rigorously defined via schema and templated SQL. The initial prototype was primarily coded in R, which was migrated to Python for enhanced functionality and integration options, e.g., visualization tools. The code and documentation are publicly accessible at https://github.com/unmtransinfo/ProteinGraphML (new Python package) and https://github.com/unmtransinfo/metap (R code). A snapshot of the currently used dataset for building **KGs** is available at http://pasilla.health.unm.edu/x/metap-pgdump.sql.gz.

**Table 1.** Data resources used to inform the development of ML-ready knowledge graphs

| **Data Source** | **Data Type** | **Data Points** |
| --- | --- | --- |
| CCLE[25] | Gene expression | 19,006,134 |
| GTEx[26] | Gene expression | 2,612,227 |
| HPA[27] | Gene & Protein expression | 949,199 |
| Reactome[28] | Biological pathways | 303,681 |
| KEGG[29] | Biological pathways | 27,683 |
| STRING[30] | Protein-Protein interactions | 5,080,023 |
| GO[31] | Biological pathways & Gene function | 434,317 |
| InterPro[32] | Protein structure and function | 467,163 |
| ClinVar[33] | Human Gene - Disease/Phenotype associations | 881,357 |
| GWAS Catalog[34] | Gene - Disease/Phenotype associations | 54,360 |
| OMIM[35] | Human Gene - Disease/Phenotype associations | 25,557 |
| UniProt[36] Disease | Human Gene - Disease/Phenotype associations | 5,365 |
| DISEASES[37] | Gene - Disease associations from text mining | 44,829 |
| NCBI Homology [38] | Homology mapping of human/mouse/rat genes | 70,922 |
| IMPC[39] | Mouse Gene - Phenotype associations | 2,153,999 |
| RGD[40] | Rat Gene - Phenotype associations | 117,606 |
| LINCS[41] | Drug induced gene signatures | 230,111,315 |

### Meta Path/XGBoost using Knowledge Graphs

The majority of biological system networks (BSN) are heterogeneous with multiple node and edge types, as illustrated by the Data Types in **Table 1**. Recent developments in heterogeneous[42,43] and BSN [44] relationship predictions introduced and formalized a new framework that takes into account BSN heterogeneity by defining type specific node-edge paths or Meta Paths[45]. A meta path [42,46] is a path consisting of a sequence of relations defined between different object types (i.e., structural paths at the meta level). In biology, the Meta Path approach can be used to seek different network paths that connect proteins / genes to specific properties such as phenotype or disease [45]. In this paper, we distinguish Meta Path (the method) from “meta paths”, used to evaluate network topology. The objects in question are graph **Nodes**, which can be genes or proteins; diseases or phenotypes; chemical structures or drugs; and other entities relevant to modeling biomedical processes. “Structural paths” reflect relationships between the different entities, i.e., Nodes. “Paths”, or Edges in topological terms, can be expression data; pathways; LINCS genomic signatures; protein-protein interactions (PPIs); or other relationships of biological relevance. Enumeration of all network paths along a defined meta path are used to compute graph topological features such as path counts by counting all path instances along that meta path. Degree weighted path counts (DWPCs — see **Eq. 1**) [44] use the number of Edges connecting each Node along a meta path to assign different weights to each path instance. Specifically, DWPCs quantify meta path prevalence using a dampening exponent (w, set to 0.4) to down weight paths through high-degree nodes when computing DWPCs:

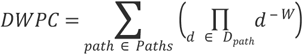

where ⊓ is the path-degree product calculated by: 1) extracting all edge-specific degrees along the path *D_path_*, where each edge contributes two degrees; 2) raising each degree *d* to the *-w* power, where *w* is the dampening exponent; 3) multiplying all exponentiated degrees to yield ⊓. DWPC is the sum of path degree products ⊓ (the DWPC section was adapted from [44]). The Meta Path formalism enables the transformation of the heterogeneous KG data types into an ML-ready input. The algorithmic workflow of applying ML to the Meta Path methodology is depicted in **Fig. 1**..

**Figure 1.**
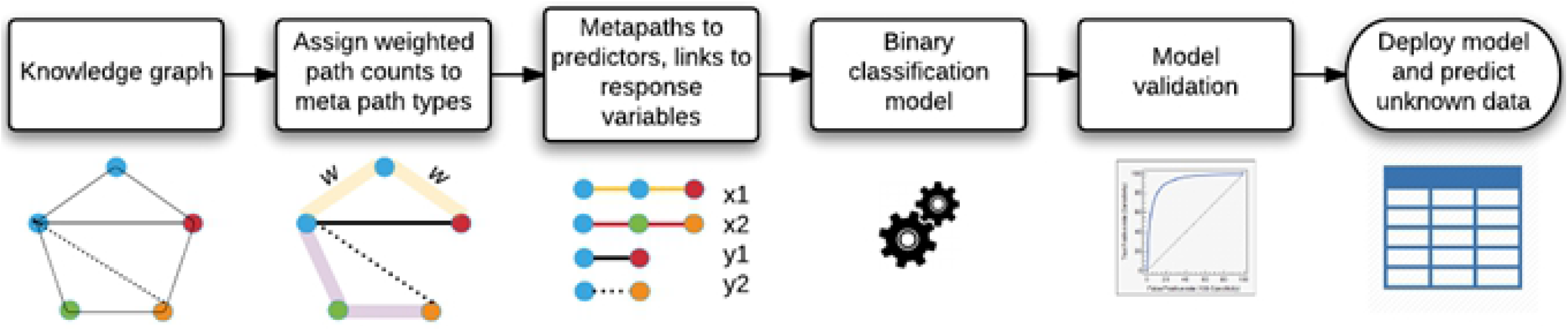
Analytical algorithms workflow for ranking target gene lists.

To better understand and predict “function” or “role in disease/phenotype”, we selected XGBoost [47], an award-winning ML algorithm that can be directly deployed onto typed meta paths. XGBoost classifiers have certain advantages compared to other ML algorithms: *i)* very fast model training and prediction, and high scalability to very large datasets; *ii)* variable selection is driven by model performance as opposed to random, such as that used in comparable ML algorithms; *iii)* built-in model interpretability. XGBoost outputs variable importance in projection (VIP) estimates, unlike neural networks and other deep learning methods. Using VIP, meta-paths are sorted in decreasing order of importance, which leads to directly interpretable insights into mechanistic processes and **Node** interactions that contribute to protein/gene “function” or “role in disease.” The combination of Meta Path/XGBoost (herein referred to as “MPxgb”) processes assertions/evidence chains of heterogeneous biological data types and identifies similar assertions. These determine the strength of the evidence linking a gene/protein to a disease or phenotype or, in this instance, autophagy.

## Development of an ATG-specific MPxgb Model

We extracted data from 17 protein-centric datasets available collected the TargetCentral relational database, TCRD [48], and summarized in **Table 1**. Specifically for this study, we started with 288 human genes from a specialized resource, the Autophagy database, ATGdb [49], which were processed and sorted for uniqueness. These genes were further queried for further confirmation of their ATG association using the “hsa04140” keyword in the Kyoto Encyclopedia of Genes and Genomes, KEGG[29] Pathway; using the gene ontology (GO) [31] term “GO:0061919”; or the UniProt [36] keyword “autophagy”, respectively. The secondary query, used to confirm the importance of specific genes in autophagy, resulted in N = 103 genes (Supplementary Material).

We trained the MPxgb model using these 103 genes as “positive” labels, i.e., known to be associated with ATG. A separate set of N = 3,468 OMIM genes, which are associated with a variety of monogenic diseases and are not present in the ATGdb, were assumed to lack ATG involvement. We generated 100 randomized MPxgb models, using 10-fold cross-validation; the median area under the curve (AUC) for these models was 0.994 ± 0.008. The model with the highest AUC (0.999932263) was selected to predict novel ATG genes, and is discussed here. The Excel file with the complete training set (3,571 genes, both positive and negative labels) is available as Supplementary Material.

## Results and Discussion

This paper introduces an autophagy-dedicated machine learning model, which is intended to serve as guidance for extending the current autophagy knowledge to gene sets that may have not been previously explored. Its main purpose is to combine current levels of evidence, and show how such evidence can be captured and modeled using a binary classifier based on the Meta Path (**Figure 2**) and XGBoost approach, built upon 17 different data sources (**Table 1**). Biologically, this model implicates a set of up to 193 previously untested genes, some of that may have a functional role in autophagy.

**Figure 2.**
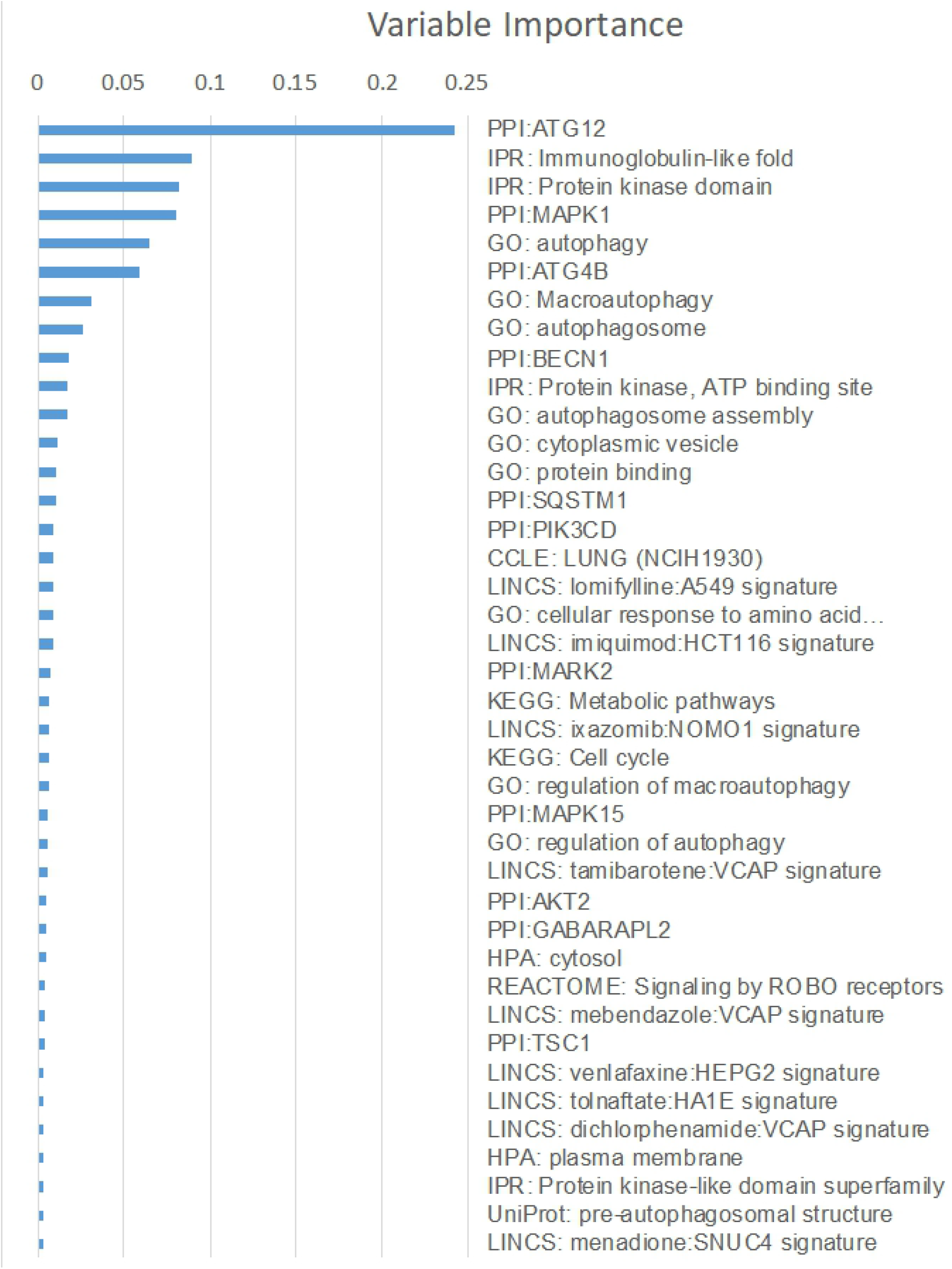
Top 40 variables, selected by the ATG MPxgb model, in decreasing order of importance. PPI: protein-protein interaction nodes; IPR: InterPro domain; GO: Gene Ontology term; HPA: Human Protein Atlas. See text for additional details.

For this model, data for 103 ATG-associated genes were contrasted with similar data for 3,468 “negative” genes, extracted from seven distinct categories of evidence: *gene & protein expression* (CCLE, the cancer cell line encyclopedia [25]; GTEx, Gene Tissue Expression [26]; and the Human Protein Atlas [27]); *biological pathways and gene function* (Reactome [28], KEGG [29] and GO [31]); *Protein-Protein Interactions* (STRING [30]); *protein structure and function* (InterPro [32]); *human gene - disease/phenotype associations* (ClinVar [33], GWAS Catalog [34], OMIM [35], UniProt Disease [36], and DISEASES [37]); *animal gene - phenotype associations* (mouse data from IMPC [39] and rat data from RGD [40], mapped to human genes via the NCBI Homology [38] resource); and *drug-induced gene signatures from the Library of Integrated Network-Based Cellular Signatures, LINCS* [41], respectively.

From these resources, meta paths were assembled, such as: {Protein — (member of) → STRING PPI Network ← (member of) — Protein — (associated with) → Disease} and {Protein — (expressed in) → Tissue/Cell line ← (associate with) — GO Term}. The first BSN application [44] to model novel disease associations using meta paths used logistic regression and ridge logistic regression. We adapted the Meta Path framework to XGBoost, a tree-boosting classification algorithm [47] used in 17 of the 29 winning solutions in Kaggle challenge competitions [50]. Combining XGBoost, a scalable ML algorithm capable of handling sparse data with BSN typed meta-paths; we used the MPxgb approach to predict novel protein functions given a variety of (unrelated) data sources.

The methodology introduced in this work has substantial advantages over simply applying ML to an existing, static dataset. **KG** construction is informed by expertise spanning several biomedical domains, with automation tools designed for an online learning paradigm to incorporate the latest findings. Predictions presented here are not generated *ex nihilo* but from the rigorously aggregated findings of autophagy research to date. Our methodology uses powerful ML tools to abstract that collective intelligence and is subject to change, and improvement as the field continues to advance.

### Model Validation through Variable Importance Selection

The top 40 features (total gain, 0.9) used to derive the best MPxgb ATG model are depicted in **Figure 2**, and available as Supplementary Material. There are 11 protein-protein interactions (PPIs), with a cumulative gain of 0.445; 4 interpro domain, IPR terms and one UniProt term (cumulative gain, 0.195); 9 GO terms (cumulative gain, 0.182); 9 LINCS terms (cumulative gain, 0.044); 3 pathway terms (cumulative gain, 0.017); and 3 expression terms (cumulative gain, 0.016), respectively. Using PPIs and structural terms provides a 64% contribution to the ATG MPxgb model; by contrast, pathway and expression terms. The presence of “autophagy” in 6 GO terms and one UniProt term provides a cumulative gain of 0.154. The related terms, “autophagy of mitochondrion”; “autophagy of nucleus”; and “autophagosome membrane”, respectively, are not present in the top 60 descriptors and, with a cumulative gain of 0.003, are not contributing to the model. This, most likely, has to do with lack of association between these terms and the 103 ATG-positive genes. LINCS-associated terms do not provide a significant contribution to the model. This is not surprising, since the LINCS descriptors used here are derived from measuring cancer cell genomic response to drug perturbation, which is perhaps less relevant in the context of autophagy. Among the top 40 descriptors are 11 PPI nodes: *ATG12, MAPK1, ATG4B, BECN1, SQSTM1*, *PIK3CD, MARK2, MAPK15, AKT2, GABARAPL2,* and *TSC1,* respectively, with the top 4 (*ATG12*, *MAPK1, ATG4B,* and *BECN1)* accounting for 40% of the total gain. Except for *MARK2* and *MAPK15*, these form a complex network of protein interactions (https://bit.ly/2JHygyq). The vast majority of the STRING [30] analysis enrichment terms point to autophagy (including macroautophagy, autophagy of mitochondrion, autophagosome assembly); phosphotransferase and kinase activity; as well as five publications related to autophagy [51–55].

Of the two outlier PPI nodes, *MARK2* and *MAPK15, MAPK15* is part of the ULK complex [56] and stimulates autophagy by interacting with ATG8 family proteins [57]. The involvement of *MARK2* in autophagy is less clear [58]. When combined with the 103 ATG-model input genes, a complex regulatory network enriched in the same autophagy terms is observed (https://bit.ly/2Z2SF60), with 3 additional KEGG pathways: *“hsa04211: Longevity regulating pathway”, “hsa04150: mTOR signaling pathway”* and *“hsa04068: FoxO signaling pathway”,* respectively. The top five publications are also related to autophagy [52,59–62]. Based on the STRING analysis of MPxgb selected PPI nodes, we conclude that these descriptors are relevant for autophagy. The MPxgb variable selection of IPR domains, as well as GO terms, is also related to genes that play a relevant role in autophagy (see also Supplementary Material). We conclude that the ATG MPxgb model discussed herein is significantly enriched in autophagy related terms and PPI nodes, and bears relevance for guiding future autophagy research with respect to potentially novel gene annotation for ATG involvement.

### Model Output: Quantifying Knowledge

We examined the top 251 predicted genes, as ranked by the predicted probability of association with ATG. Among these, 9 were present in the initial set of 288 ATGdb genes; another 34 genes are annotated in the ATGdb expanded set (see Supplementary Material). A total of 43 genes predicted by the ATG MPxgb model were retrieved from ATGdb. Another set of 15 genes could be found by performing additional queries, such as “Autophagy Pathway” in PathCards [63]. Thus, 58 of the top 251 genes, or 23.1%, appear to have confirmed (or strongly suspected) association with autophagy. However, 193 of these top 251 genes do not appear to be associated with autophagy, despite efforts to consult a variety of literature-based resources and online databases (which include ATGdb, PathCards, KEGG, GO, UniProt and TCRD).

Given that most of the top 40 ATG MPxgb variables and 23% of the top 251 genes (including the gene ranked 246, *ARFGEF1* - present in ATGdb), we posit that this machine learning model bears relevance in the study of autophagy, and suggest that some of these 193 genes may represent autophagy “dark genes”. We also examined the “Target development level” (TDL), a knowledge-based classification for human proteins [22] that can be used to explore the dark genome [64]. In brief, **Tclin** are proteins via which approved drugs act (i.e., mode-of-action drug targets); **Tchem** are proteins known to bind small molecules with high potency; **Tbio** are proteins with well-studied biology, having a fractional publication count above 5 [65] or well-annotated OMIM (disease) phenotypes; and **Tdark** are understudied proteins that do not meet criteria for the above 3 categories, respectively. The TDL count for the training set (103 ATG-associated genes), the test set (top 251 predicted genes) and the “ATG dark genes?” subset (193 genes) is summarized in **Table 2**. As estimated by this knowledge classification system, it can be concluded that the majority of the predicted genes, specifically for the ATG dark genes, are either Tbio or Tdark. Indeed, nearly 30.6% of the “ATG dark genes” are classified as **Tdark**. which suggests that a significant portion of these genes are understudied.

**Table 2.** Target development level distribution for the ATG MPxgb model sets

| Set | Tclin | Tchem | Tbio | Tdark |
| --- | --- | --- | --- | --- |
| Training | 10 | 32 | 60 | 1 |
| Test | 0 | 52 | 137 | 62 |
| ATG dark gene? | 0 | 37 | 97 | 59 |

### Model Output: Newly Predicted Genes

For the sake of brevity, we focus the remainder of this discussion on the top 40 predicted genes, which are outlined in **Table 3**. Of these, 15 genes are already in ATGdb (*GABARAPL3, ULK3, EIF2AK2, RRAGD, RAB1A, VPS39, CSNK1G2, SNX1, CSNK2A2, PRKAB1, RAB9A, PRKCI, MAP3K14, EXOC4, CALCOCO2). RMDN1,* interacts with Beclin 1, a known regulator of autophagy [66]; another gene, SLK, escorts *VPS4B* which itself is involved in autophagy [67]; *OXSR1* is upstream of AMPK, a known autophagy regulator [68]; whereas *MTMR3* and *BNIP3L* are ATG-annotated in PathCards.

**Table 3.**
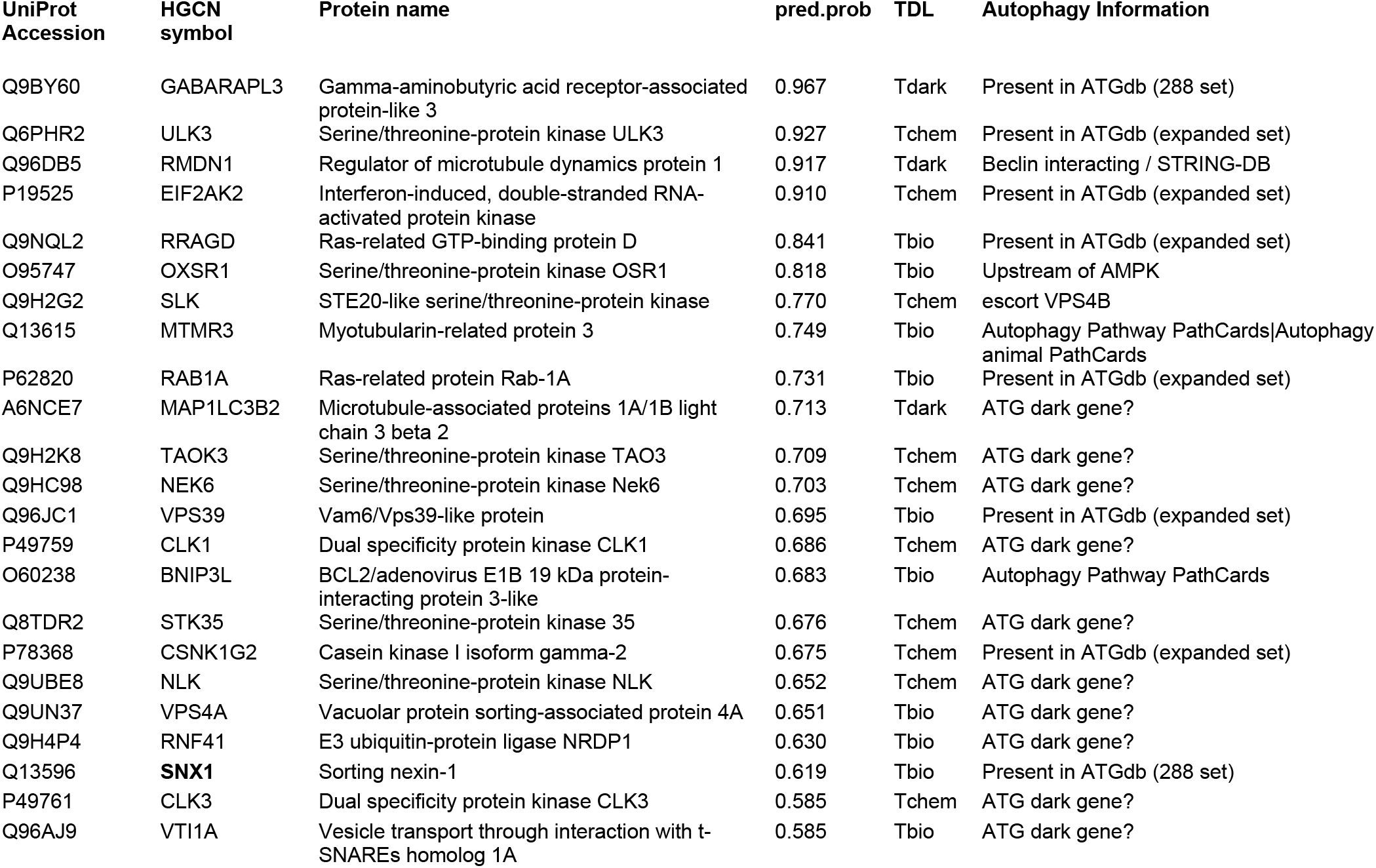
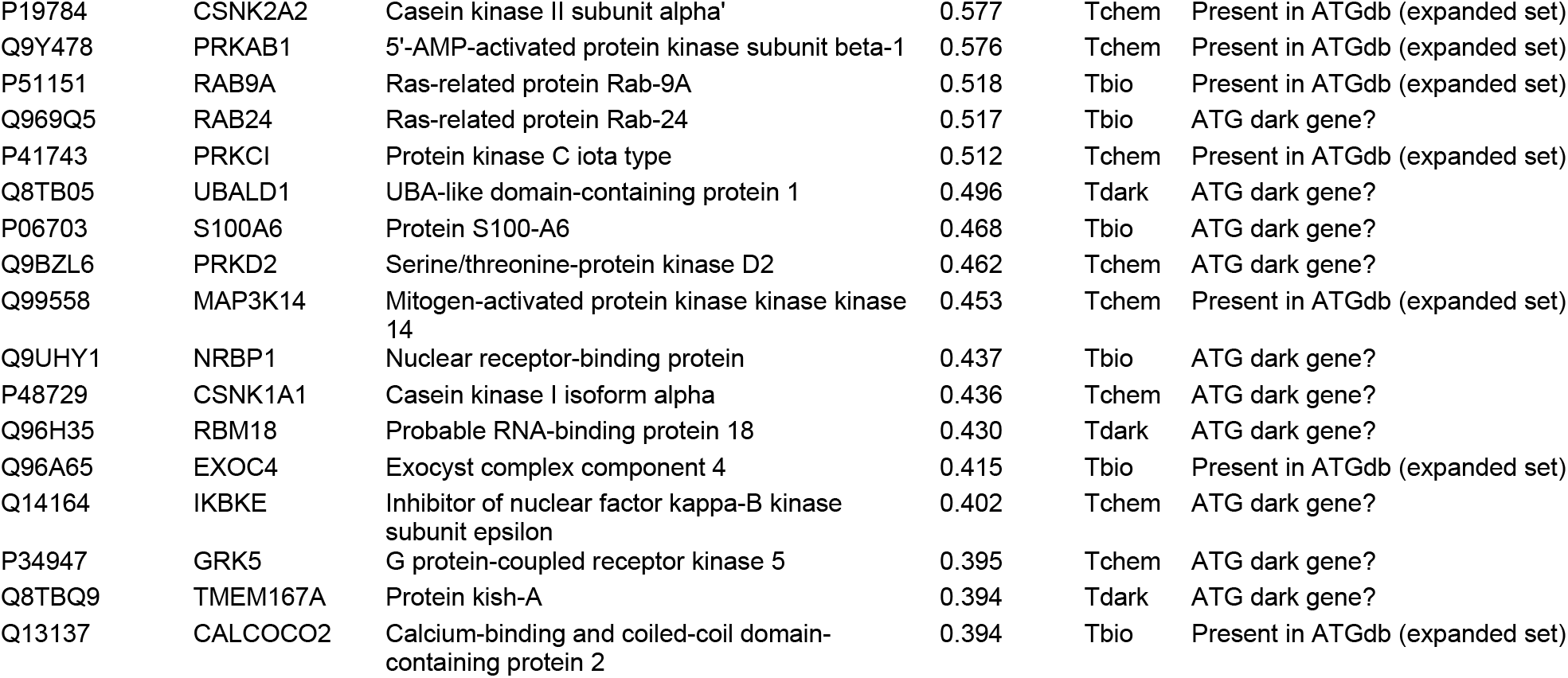
List of top 40 genes from the ATG MPxgb model. HUGO Gene Nomenclature Committee. Pred.prob - predicted probability, as derived from the model.

Twenty of the top 40 genes may represent, however, novel assertions with respect to autophagy: *MAP1LC3B2, TAOK3, NEK6, CLK1, STK35, NLK, VPS4A, RNF41, CLK3, VTI1A, RAB24, UBALD1, S100A6, PRKD2, NRBP1, CSNK1A1, RBM18, IKBKE, GRK5,* and *TMEM167A,* respectively. With six exceptions *(TAOK3, CLK1, CLK3, UBALD1, RBM18,* and *TMEM167A*), the remaining 14 genes form a complex network when combined with the 11 PPI nodes discussed in the Variable Importance section (https://bit.ly/2Yc5EoZ). As previously mentioned, the STRING analysis enrichment terms point to autophagy (including autophagy, macroautophagy, and autophagosome assembly); protein serine/threonine kinase activity activity; KEGG Pathway terms *“hsa04140: Autophagy - animal”, “hsa05167: Kaposi’s sarcoma-associated herpesvirus infection”, “hsa05160: Hepatitis C”* and *“hsa04152: AMPK signaling pathway”* as well as multiple publications related to autophagy [52,53,55,60,69–72].

Since many of the potentially novel 20 genes out of the top 40 are kinases, we used X2K Web (eXpression2Kinases) [73] to perform a Transcription Factor Enrichment Analysis (TFEA) [74]. This type of analysis predicts which transcription factors (TFs) are most likely to regulate the expression of these 20 genes. The ranked list of predicted TFs is visually summarized in **Figure 3**. The top ranked 13 transcription factors, selected to ensure that each of the top 20 predicted autophagy genes is represented at least once, are as follows: *RUNX1, TCF7L2, ELF1, TAF1, GATA1, FOXP2, FOXA1, SPI1, CHD1, SRF, FOXA2, ZNF384,* and *BRCA1,* respectively.

**Figure 3.**
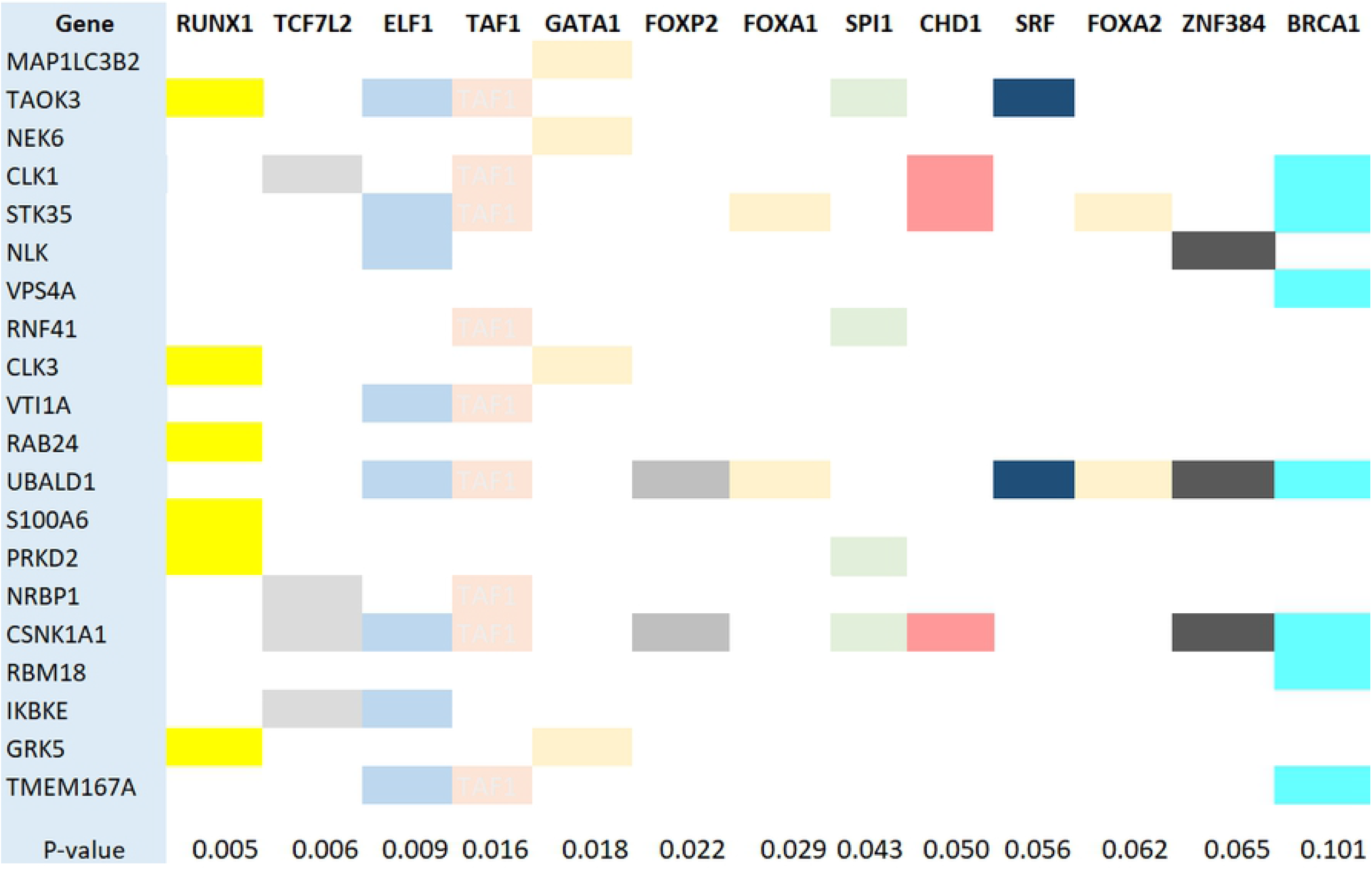
Visual summary of the transcription factor enrichment analysis for the top 20 predicted genes autophagy dark genes. Thirteen transcription factors are shown in reverse order of P-value ranking. See text for additional details.

When combining the 11 PPI node genes selected by the ATG MPxgb model with the 13 genes TFEA-selected genes, the top potentially novel ATG genes form a complex STRING-based network (https://bit.ly/2YeOffr) that leaves six genes out *(NRBP1, CLK1, CLK3, UBALD1, RBM18*, and *TMEM167A)*. In addition to GO terms related to autophagy, the STRING analysis enrichment terms include “transcription factor binding” and “membrane-bounded organelle”; the UniProt keyword “Phosphoprotein”; the KEGG Pathway terms *“hsa03022: Basal transcription factors”, “hsa04137: Mitophagy - animal”* and *“hsa05168: Herpes simplex infection”,* as well as several publications related to autophagy [51,52,55,69,75–77]. While the TFEA step does appear to include STRING enrichment terms associated to TFs, it also provides additional data elements supporting the role of some proteins left out of the STRING-based PPI network: *UBALD1* might be regulated by 10 of the 13 TFs, *TAOK3* and *CLK1* by 5, and *TMEM167A* by 4 TFs, whereas *NRBP1* and *RBM18* are regulated by 2. Both this and the prior STRING network highlight the relationship between some ATG-related genes and viral *(H. simplex*, *Hepatitis C)* infection. This is not surprising, since autophagy is an antiviral defense mechanism.

## Conclusions

In this paper we introduced an ATG-dedicated machine learning model, which we intended to provide as guidance for extending current autophagy knowledge, in order to explore gene that previously have not been evaluated for their role in autophagy. In developing this ATG-specific MPxgb model, we started with 288 genes associated with autophagy from the Autophagy database, ATGdb [49], upon which we set additional filters (association with GO, KEGG and UniProt terms), to increase confidence in the remaining 103 positive label genes. As negative labels, we used N = 3,468 genes associated with monogenic diseases in OMIM, which were not present in ATGdb, as genes lacking the ATG association. This assumption, which is necessary for many machine learning methods, specifically for binary classifiers, may represent a significant flaw since we lack absolute certainty that negative examples are *not* genes playing a role in autophagy. However, just as any other ML models, the ATG-specific MPxgb model requires extensive validation, perhaps via several iterations. Reflecting current levels of evidence, we used XGBoost, a binary classifier combined with the Metapath (**Figure 2**) approach, which was deployed upon 17 sources of data representing seven distinct categories (**Table 1**). The ATG MPxgb model implicates up to 193 previously untested genes that may have a functional role in autophagy. This was, in fact, our primary motivation: *to provide the scientific community with a ML-ready list of autophagy genes, as summarized by the training set, combined with a truly blind prediction set (N = 193)*. By disclosing this list of putative genes, we aim to encourage further experimental testing. All gene sets mentioned throughout this paper are available in the Supplementary Material. Future validation steps may include integration of chemical–protein annotation resources, using Pharos [48] and Chem-Prot [78], the use of a semantic model to evaluate druggability via the drug target ontology [79].

## Acknowledgments

This work was supported by NIH grants U24 CA224370 (TIO, DB, JJY), U24 TR002278 (TIO) and P20 GM121176 (TIO, VD). Dr. Oleg Ursu (Merck Research Laboratories) implemented the initial version of the MPxgb algorithm.

